# Remembering touch: Somatosensory cortex supports long-term memory for dynamic vibrotactile patterns

**DOI:** 10.64898/2026.09.27.754826

**Authors:** Marianna Elisa Schmidt, Timo Torsten Schmidt, Till Nierhaus, Felix Blankenburg

## Abstract

The neural substrates of tactile long-term memory (LTM) in humans remain poorly understood, as relatively few neuroimaging studies have addressed this question. Here, we asked whether retrieval of tactile stimulation engages modality-specific representations in the primary and secondary somatosensory cortices (S1 and S2), alongside domain-general retrieval networks. Participants learned 16 tactile trajectories, each presented for 1.2 s on the left thumb through sequential activation of 10 vibrating dots on a Braille-like display. Three learning sessions within one week were followed by a recognition test during functional magnetic resonance imaging (fMRI). In the final sample (n = 26), we contrasted BOLD responses during successful recognition of learned stimuli with those during correct rejection of novel stimuli. Successful recognition was associated with increased activity in contralateral S1 and S2, alongside activation in the hippocampus as well as frontal and parietal regions implicated in memory retrieval. A psychophysiological interaction analysis (PPI) further revealed increased task-dependent connectivity between the hippocampus and contralateral S1 and S2 during successful recognition. Together, these findings suggest that tactile long-term memory retrieval relies on coordinated hippocampal-somatosensory interactions and may engage long-term memory representations in somatosensory cortex.

## 1 Introduction

Remember the feeling of sand beneath your feet, slipping between your toes the last time you stood on a beach. For a moment, it can feel as if you are experiencing such a tactile event again. This vividness is thought to arise, at least in part, from the reinstatement of neural activity patterns engaged during the original sensory experience (Wheeler et al., 2000). In this view, long-term memory representations, often conceptualized in terms of memory engrams (Josselyn & Tonegawa, 2020; Kitamura et al., 2017), involve distributed patterns of activity spanning modality-specific sensory cortices and higher-order memory systems. The hippocampus is thought to support retrieval through pattern completion and reactivation of distributed cortical representations, while prefrontal and posterior parietal regions contribute to controlled retrieval, evaluation and integration of recovered mnemonic information (Danker & Anderson, 2010; Moscovitch et al., 2016).

While extensive work has characterized contributions of sensory systems to long-term memory (LTM) in the visual and auditory domains (Danker & Anderson, 2010), considerably less is known about neuronal representations underlying tactile LTM. Evidence from visual and auditory memory suggests a division of contributions along the sensory hierarchy, whereby primary sensory cortices represent relatively low-level stimulus features, whereas higher-order regions support more integrated or abstract representations of the memory content (Cooke et al., 2015; Gaffan, 2002; Loo et al., 2021; Mishkin, 1982; Mishkin & Appenzeller, 1987; Weinberger, 2004). S1 contains fine-grained, somatotopically organized maps of individual body parts and digits, including spatially distinct finger representations (Sakai et al., 1995; Sanchez-Panchuelo et al., 2010). S2, in contrast, is characterized by larger and more complex receptive fields and greater integration of tactile information across digits, body sides, and over time, allowing for the encoding of more complex and integrated features of tactile stimuli (Zhu et al., 2007). Both regions are recruited during tactile perception as well as internally generated tactile experiences, including tactile mental imagery (Nierhaus et al., 2023; Schmidt et al., 2014; Schmidt & Blankenburg, 2019; Schröder et al., 2019). Together, these properties suggest that S1 may contribute to the reinstatement of body-part-specific sensory features of tactile long-term memories, whereas S2 may support the reactivation of more integrated spatiotemporal features of the remembered tactile pattern.

Empirical research of tactile LTM processes in humans remains limited and has yielded inconclusive findings. Early positron emission tomography (PET) work identified a ventral parietoinsular network, including S2, insula, and medial temporal regions, during retrieval of haptically encoded novel shapes (Bonda et al., 1996). By contrast, a functional magnetic resonance imaging (fMRI) study of haptically encoded objects reported recruitment of S1 alongside inferior parietal and frontal regions, during memory retrieval (Stock et al., 2009). Together, these findings implicate both early somatosensory and higher-order associative regions in tactile LTM but leave the respective contributions of S1 and S2, as well as their interactions with hippocampal memory systems, unresolved. Moreover, previous studies largely relied on the active exploration of complex haptic objects (Bonda et al., 1996; Stock et al., 2009), in which tactile information is inherently coupled with motor and potentially multisensory processing. Together these make it difficult to isolate modality-specific tactile memory representations for ecologically valid tactile events.

Here, we address these gaps by testing the neural substrates of tactile LTM using an fMRI recognition paradigm for learned vibrotactile stimulation events. Thirty participants were trained over three consecutive days to memorize 16 distinct dynamic tactile stimuli, which resemble relatively naturalistic tactile events of a vibrating dot moving for 1.2 s along a patch of skin on the left thumb. Recognition memory was tested during fMRI, at least 24 h after training, to ensure testing long-term memory representations. We hypothesized that contralateral S1, particularly the cortical area corresponding to the stimulated thumb, and S2 contributing to modality-specific tactile components of distributed tactile memory engrams, whereas hippocampal, frontal and parietal regions support broader retrieval processes. We further tested whether successful tactile memory retrieval is accompanied by increased functional coupling between hippocampal memory systems and somatosensory cortices. Together, these analyses allowed us to examine how modality-specific somatosensory representations interact with distributed memory systems during long-term tactile memory retrieval.

## 2 Methods

### 2.1 Participants

Thirty participants without neurological or psychiatric disorders completed the study procedures after giving written informed consent. Four datasets were excluded due to insufficient task performance (see Results) or excessive head motion, predefined as more than 5% of all volumes exceeded a frame-wise displacement (FWD) threshold of 0.4 mm (Power et al., 2012). The final sample comprised n = 26 participants (mean age in years: 25.9 ± 5.3; 15 females; 11 males; 4 left-handed; 22 right-handed), whose data entered the main analyses. The study adhered to the Human Subject Guidelines of the Declaration of Helsinki and was approved by the local ethics committee of Freie Universität Berlin (003/2021, Berlin, Germany).

### 2.2 Experimental stimuli

Stimuli were designed to evoke the sensation of a vibrating point moving across the thumb, requiring the integration of successive tactile events into a coherent spatiotemporal stimulation pattern. This sequential structure was chosen to engage temporal binding processes associated with hippocampal memory formation (Eichenbaum, 2013; Paz et al., 2010; Tubridy & Davachi, 2011), while approximating the temporally extended nature of natural tactile experiences under highly controlled experimental conditions. An example stimulus configuration is shown in Figure 1A. Vibrotactile stimuli were delivered using a 32-dot piezoelectric Braille-like display (4 x 8 matrix with 2.5 mm spacing) attached to the left thumb, and controlled by a programmable stimulator (Piezostimulator, QuaeroSys, St. Johann, Germany). Each stimulus presentation lasted 1.2 s and consisted of a sequence of vibrating pins across 10 consecutive positions on the display, creating the sensation of a single vibrating point moving across the thumb. Each position was activated only once within a given stimulus sequence. To create the sensation of smooth, continuous movement, activation of each successive pin began slightly before the one of the previous pin ended.

**FIGURE 1.**
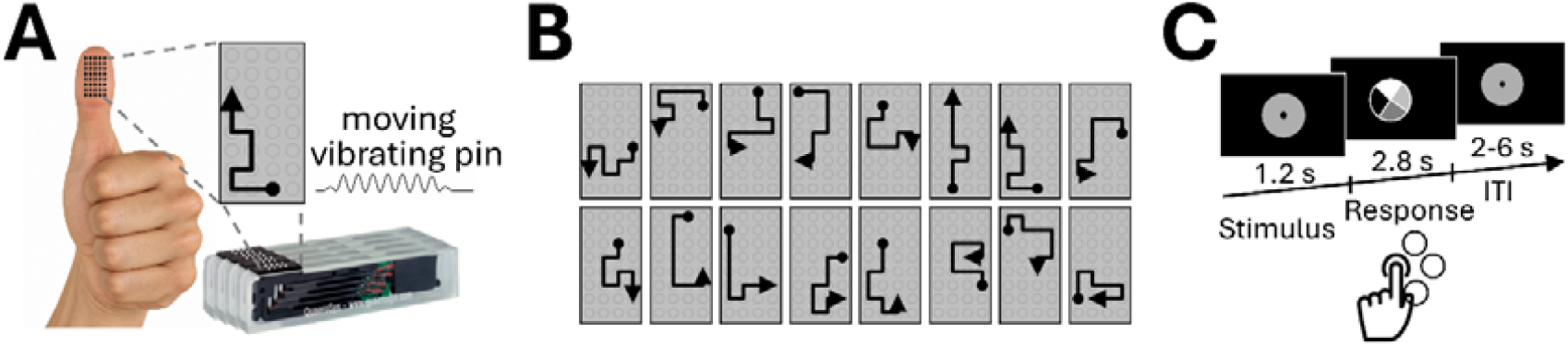
Experimental design. (A) Stimuli were presented as dynamic smooth movement of a vibrating pin across a trajectory on a 4 × 8 pin display. Each stimulus comprised a sequence of 10 pin positions. Each stimulus lasted 1.2 sec for the pin to move through the display. (B) A full example stimulus set of one participant is illustrated. (C) Recognition task during fMRI. Each trial consisted of one stimulus presentation, followed by a 2.8 sec response period and a variable ITI (2–6 sec). Participants classified each stimulus as learned or novel and simultaneously indicated response confidence using four response categories: sure learned, unsure learned, unsure novel, and sure novel. The mapping of these response categories onto the four buttons of a diamond-shaped response box varied across trials and was indicated by the display during the response window. Each fMRI run comprised 48 trials, and six runs were acquired. For the fMRI session, novel stimuli were generated by reversing the pin sequence of learned stimuli while preserving their spatiotemporal characteristics, whereas in the learning session newly generated foils were used.

For each participant, an individual set of 16 to-be-learned stimuli was generated (Figure 1B). Participant-specific stimulus sets were used to minimize systematic item effects across participants, such that differences in the intrinsic discriminability or memorability of individual trajectories would not consistently bias behavioral performance or neural responses. Stimuli for each set were sampled from all possible trajectories across the display, with each path lengths comprising ten pin positions. To ensure sufficient dissimilarity among stimuli within a set, the spatial trajectories of any two stimuli were required not to exceed a correlation of ρ = 0.50. During the recognition tests in the learning sessions (see below), additional randomly chosen stimuli were used as foil. For the fMRI recognition test, the 16 learned stimuli were presented together with 16 novel stimuli generated by reversing the path direction of the corresponding learned trajectories. This was done to match the physical stimulus properties of learned and novel stimuli in the fMRI part as closely as possible to ensure that during stimulation no systematic differences in the afferent cortical area in S1 was created. Participants were not exposed to the reversed versions before the scanning session, ensuring that these stimuli were novel at the time of the fMRI recognition test.

### 2.3 Learning sessions

Participants were invited to four experimental sessions within a seven-day period, with learning sessions conducted either on consecutive days or separated by one or two days. The first three appointments were learning sessions, whereas the fourth was the fMRI recognition test.

During each learning session, participants were trained in six learning runs, each followed by a recognition test. Across each session, participants were trained on all 16 stimuli of their individual stimulus set. During the first three runs, participants were trained on half of the stimulus set (eight stimuli), whereas the remaining eight stimuli were trained during the second three runs. This subdivision was used to reduce the number of stimuli learned concurrently and thereby facilitate acquisition of the individual tactile patterns. Assignment of stimuli to the first or second half was balanced across the learning sessions. Each learning run comprised five stimulation phases of 1 min 40 s. Within each phase, each of the eight to-be-learned stimuli was presented three consecutive times. The presentation order of the eight stimuli was held constant for the five stimulation phases within a given run (e.g., 3,3,3,7,7,7,2,2,2,8,8,8, …) but was randomized between runs. Stimuli were separated by intertrial intervals (ITIs) of 2, 3 or 4 s.

Participants were informed that they needed to learn a total of 16 stimuli and that eight stimuli would be presented during each half of the learning session. They were further informed that the stimuli do not come with a specific label or order. Participants were instructed to memorize all presented stimuli while focusing on their tactile characteristics to minimize the use of alternative learning strategies. During stimulation, participants kept their eyes open and fixated a central cross while wearing passive noise-attenuating headphones to eliminate auditory cues from the stimulation device. No explicit task was required during stimulus presentation other than memorization. At the end of each learning run, participants completed a brief recognition test using the same task as in the fMRI experiment. Each test comprised the eight trained stimuli and eight randomly selected novel stimuli, presented across 16 trials and lasting approximately 2 min (see below). Additionally, at the end of each learning session a recognition test on all 16 stimuli like the one during the fMRI session, but with randomly chosen novel stimuli, was conducted to monitor the learning progress.

### 2.4 Recognition test during fMRI

During fMRI, participants completed six runs of the recognition task, each comprising 48 trials and lasting approximately 7 min per run (Figure 1C). In each run, all 16 learned stimuli were presented once, with eight of these stimuli presented a second time, resulting in 24 learned-stimulus trials. These were accompanied by 24 novel-stimulus trials, yielding an equal number of learned and novel trials per run. In every trial, participants were presented with one stimulus. The task for the participants was to report if that stimulus was learned or novel (not presented before). Only in the fMRI recognition task, novel stimuli were generated from the learned stimuli by presenting the trajectories of the learned stimuli in reverse temporal order, thereby closely matching the physical and somatotopic properties of learned and novel stimuli while differing in their spatiotemporal sequence. Participants chose one of four response options: (1) sure learned, (2) unsure learned, (3) unsure novel, (4) sure novel. The four response buttons were arranged in a diamond-shape on the button-box. To prevent systematic effects of motor preparations the mapping of response option to response buttons varied for every trial. Following stimulus presentation, a diamond-shaped visual response cue indicated the current response mapping using four different shades of gray corresponding to the four response options. The mapping between gray levels and response categories was identical across participants. The response was given by button press using the right thumb during the display of the response cue, which shortly flickered upon response but stayed on until the end of the 2.8 s response window. During the recognition tests in the learning sessions, feedback was provided at the end of each trial (i.e., if a response was correct or incorrect). No trial-wise feedback was given during scanning to minimize additional learning effects. The ITI varied between 2, 3, 4, 5, and 6 s. At the end of each recognition test, participants were informed about their overall performance.

Recognition responses were classified according to signal detection theory as Hits (learned stimuli correctly recognized as learned), Misses (learned stimuli incorrectly judged as novel), Correct Rejections (CRs; novel stimuli correctly identified as novel), and False Alarms (FAs; novel stimuli incorrectly judged as learned), pooling across confidence levels. Recognition accuracy during the learning sessions and fMRI recognition test was calculated as the proportion of correct responses (Hits + CRs) across all trials and expressed as a percentage. Two separate repeated-measures ANOVAs tested for differences in performance across learning sessions and across fMRI runs. For supplementary behavioral analyses, the proportion of trials classified as Hits, CRs, FAs, and Misses was calculated for each participant. To assess whether repeated presentation of learned stimuli within an fMRI run affected recognition performance, an additional repeated-measures ANOVA compared performance between the first and second presentation of the stimuli that were repeated within runs. Response times (RTs) were averaged across trials and runs separately for each recognition outcome and compared using a repeated-measures ANOVA with recognition outcome (Hit, CR, FA, Miss) as the within-subject factor. To assess whether recognition conditions differed systematically in motor responses, the relative use of each of the four physical response buttons was calculated separately for each recognition outcome. Finally, the distribution of confidence responses (sure learned, unsure learned, unsure novel, sure novel) was examined across participants. When a significant main effect was observed, post hoc pairwise *t*-tests were performed with Bonferroni correction for multiple comparisons.

Although participants were instructed to focus on the tactile characteristics of the stimuli during learning, they may have additionally employed other memorization strategies. Thus, after the fMRI scanning the participants completed a debriefing questionnaire about their learning strategies which included open questions like “*What did you do to memorize the patterns*?” and ratings where the participants had to indicate on a 5-point Likert scale in how far they have used a visual, spatial, tactile, or verbalized learning strategy (1 = not at all, 5 = very strongly).

### 2.5 MRI data acquisition

fMRI data was acquired in 6 runs of 7 min each on a 3 T TIM Trio (Siemens) at the Center for Cognitive Neuroscience Berlin (CCNB). For each run, 210 functional volumes comprising 37 slices were acquired in ascending order using T2*-weighted gradient-echo EPI with a repetition time (TR) of 2 s; echo time (TE) of 30 ms; an isotropic voxel size of 3 mm; a flip angle of 70°; and 64 x 64 matrix size. Additionally, a T1-weighted structural scan using an MPRAGE sequence with 176 sagittal slices, TR = 1.9 s, TE = 2.52 ms, and an isotropic voxel size of 1 mm was acquired.

### 2.6 FMRI data preprocessing and analysis

To check for motion during the fMRI scans, the FWD rate (Power et al., 2012) was calculated for each volume using the BRAMILA tools (https://github.com/kauttoj/bramila). Pre-processing and general linear model (GLM) analysis of the fMRI data was performed with SPM12 version v7771 (Friston et al., 2007; https://www.fil.ion.ucl.ac.uk/spm/software/spm12/). Functional data were realigned to the mean image, normalized to MNI space using unified segmentation, interpolated to an isotropic voxel size of 2 mm, and spatially smoothed with an 8 mm full width at half maximum Gaussian kernel which is recommend as default kernel size in task-based fMRI experiments (Friston et al., 2007). As a last step, the functional data was detrended using a voxel-level linear model of the global signal (Macey et al., 2004).

At the first level, a separate GLM was fitted to each participant’s fMRI data. Four regressors of interest represented the signal-detection outcomes: Hits, FAs, CRs, and Misses. To account for response-related activity, four button-specific motor regressors modeled the actual time of each button press. Six head-motion parameters were included as additional nuisance regressors. To account for trial-by-trial differences in RT (see Results), a single RT regressor comprising all trials irrespective of condition was included as a further nuisance regressor. RTs were mean-centered across all valid trials within each run and entered as parametric weights at the corresponding response window onsets. The resulting event series was convolved with the canonical hemodynamic response function. This approach modeled variance associated with trial-by-trial RT while preserving the four categorical regressors used to estimate memory-related effects. Confidence ratings (sure versus unsure) were not included in the fMRI analyses because their distributions were imbalanced in several participants, with some runs lacking observations in individual confidence categories (see results).

The primary contrast of interest compared Hits with CRs, thereby contrasting successful recognition of previously learned tactile stimuli with correct identification of novel stimuli. Individual first-level contrast images were entered into a second-level random-effects GLM to assess group-level effects. Statistical significance was determined using an initial voxel-wise cluster-defining threshold of *p* < 0.001, followed by cluster-level family-wise error (FWE) correction at *p* < 0.05.

As supplementary analyses, we first assessed whether individual differences in learning strategy influenced the primary activation pattern. The second-level Hits > CRs analysis was repeated with participants’ self-reported visual, spatial, tactile and verbal learning-strategy ratings from the post-scan debriefing questionnaire entered as mean-centered covariates of no interest. We further examined several whole-brain contrasts to characterize the effects underlying the primary Hits > CRs comparison. CRs > Hits tested for greater activity during correct rejection of novel stimuli than during successful recognition of learned stimuli. Hits > FAs compared trials associated with the same “learned” response but differing in stimulus history and accuracy, thereby testing for activity related to successful recognition beyond the recognition response itself. Conversely, Misses > CRs compared trials associated with the same “novel” response but differing in stimulus history and accuracy, testing whether prior learning modulated activity even when recognition failed. Finally, Hits > Misses directly compared successful and unsuccessful recognition of previously learned stimuli, thereby testing for activity associated with successful recognition while holding stimulus history constant.

To investigate task-dependent functional connectivity during tactile memory retrieval, we performed a psychophysiological interaction (PPI) analysis using the left hippocampus as a seed region. The left hippocampus was selected based on its activation in the group-level Hits > CRs contrast (see Results). For each participant, the six experimental runs were concatenated to obtain a single continuous time series. The search space for the individual seed locations was constrained by a hippocampal mask in MNI space comprising CA1, CA2, CA3 and the dentate gyrus, derived from the JuBrain Anatomy Toolbox (Eickhoff et al., 2005). Within this mask, the participant-specific local maximum for the Hit > CR contrast was identified, and a spherical seed region with a 4-mm radius was centered on this peak. The blood-oxygenation-level-dependent (BOLD) time series of the seed-region was summarized by the first eigenvariate across voxels within the sphere.

The PPI term represented the interaction between the physiological signal from the hippocampal seed and the psychological contrast of interest, with Hits weighted +1 and CRs weighted −1. First-level PPI models included the PPI interaction term, the hippocampal physiological regressor and the psychological Hits > CRs regressor. To reduce variance related to processes not specific to the Hits > CRs comparison, the models additionally included the common all-trial RT regressor, four button-specific motor-response regressors, separate regressors for FA and Miss trials, six head-motion parameters, and run-specific block regressors accounting for concatenation of the six runs. The first-level PPI contrast tested the PPI interaction term, and the resulting contrast images were entered into a second-level random-effects GLM to identify regions showing stronger task-dependent functional coupling with the hippocampus during successful recognition of learned tactile stimuli relative to CRs of novel stimuli.

Because our primary hypothesis concerned task-dependent coupling between the hippocampus and somatosensory cortex, the main PPI analysis was restricted to an anatomically defined mask comprising right S1 and right S2 from the JuBrain Anatomy Toolbox. Restricting the analysis to these a priori regions increased anatomical specificity and focused statistical inference on the somatosensory regions most directly relevant to the tactile retrieval hypothesis. Within this mask, effects were considered significant at p < 0.001, uncorrected. To assess the broader spatial distribution of hippocampal coupling without imposing this anatomical constraint, we performed an unmasked PPI analysis using the same statistical threshold; these results are reported in the Supplementary Material.

## 3 Results

### 3.1 Recognition performance and behavioral results

Behavioral performance across the three learning sessions is shown in Figure 2A. During the recognition test of all 16 stimuli conducted at the end of each session, participants achieved a mean accuracy of 57.9% (SD = 10.1%, range = 29–79%) in the first session, followed by 61.5% (SD = 10.9%, range = 44–79%) and 64.2% (SD = 7.3%, range = 52–79%) in the second and third learning sessions, respectively. A repeated-measures ANOVA revealed a significant effect of session on recognition performance (*F*(2, 25) = 4.18, *p* < 0.05). Bonferroni-corrected paired *t*-tests showed a significant increase in performance from the first to the third session (*t* = 3.35, *p* < 0.01).

**FIGURE 2.**
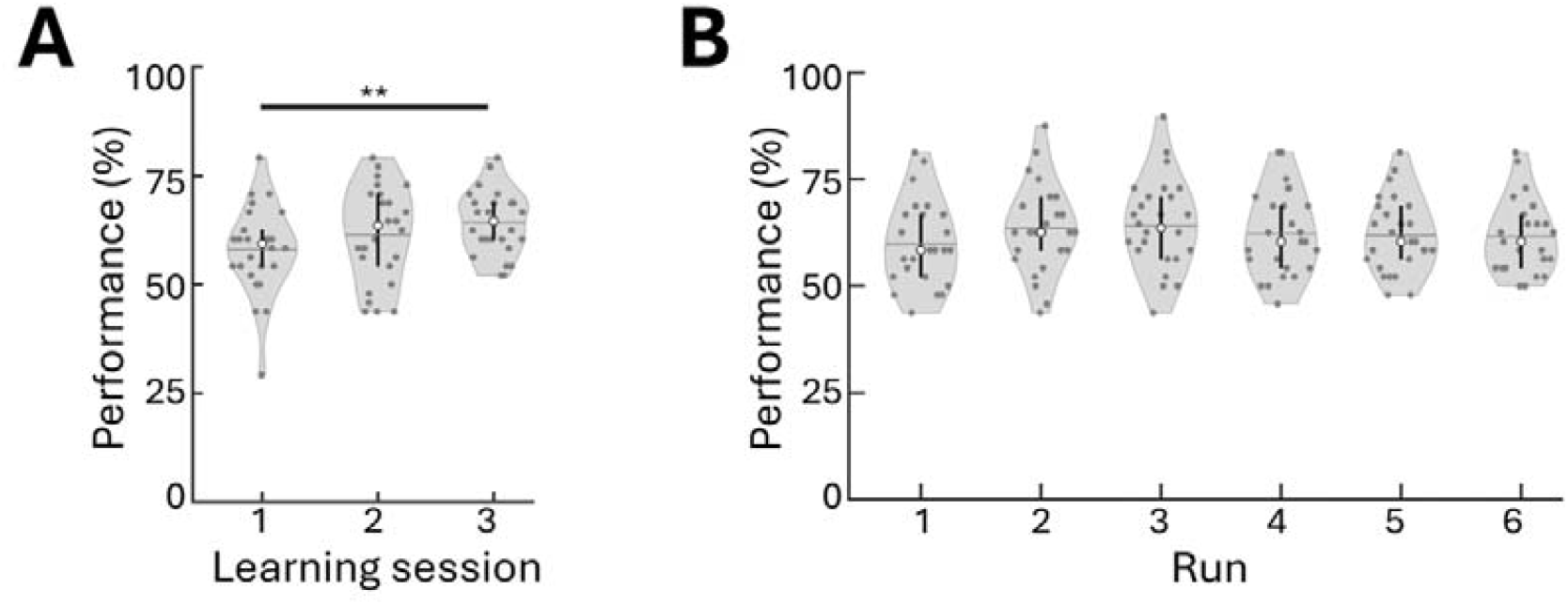
Behavioral performance during tactile long-term memory learning and recognition. (A) Recognition performance in the test administered at the end of each of the three learning sessions. Each value represents a participant’s recognition accuracy across the full set of stimuli tested at the end of that session. Performance increased from the first to the third learning session. (B) Recognition performance during the fMRI recognition test, shown separately for each of the six runs. Performance remained stable across runs, with no significant differences between runs. Violin widths represent the kernel-density estimate of participant-level performance, with wider sections indicating a higher density of observations. Gray dots indicate individual participants (n = 26), white circles indicate the median, vertical black lines span the interquartile range (25th–75th percentiles), and horizontal lines indicate the mean. Asterisks indicate significant Bonferroni-corrected pairwise comparisons following the repeated-measures ANOVA (**: *p* < 0.01).

During the fMRI recognition task, three participants showed mean recognition accuracy below chance level (<50%; 47.2%, 45.8%, and 45.8%) and were therefore excluded from further analyses. Another participant was excluded because of excessive head motion according to the predefined exclusion criterion (i.e., >5% high-motion volumes). Head motion was generally low among the included participants. On average, 1.1% of all acquired volumes exceeded the FWD threshold of 0.4 mm (SD = 1.3%, range = 0.0–6.0%).

The mean recognition accuracy during the fMRI sessions of the included n = 26 participants was 59.8%, 63.5%, 64.0%, 62.3%, 61.9% and 61.5% in run 1 to 6, respectively (Figure 2B). Overall, a mean performance of 62.8% (SD = 1.5%, range = 59.8%–64.0%) was achieved and a repeated-measures ANOVA did not reveal any significant differences in performance between runs (*F*(5, 25) = 1.9; *p* = 0.1). The distribution of signal-detection outcomes across participants is shown in Supplementary Figure 1. Across the group, Hits and CRs accounted for 32.7% and 30.1% of trials, respectively, whereas FAs and Misses accounted for 19.9% and 17.3%. RTs differed between recognition outcomes, with responses being fastest for Hits and slowest for Misses (Supplementary Figure 2A). The use of the four response buttons was comparable across recognition outcomes (Supplementary Figure 2B), indicating that differences between conditions were not systematically associated with individual motor responses. With regards to effects of repeated stimulus presentation of the learned stimuli, a repeated-measures ANOVA did not show any difference with respect to performance between the first and the second stimulus presentation within runs (*F*(1, 5) = 0.09; *p* = 0.76).

Our paradigm included a measure of recognition confidence, with participants indicating whether they were sure or unsure about each response. However, the distribution of confidence ratings varied substantially across participants (Supplementary Figure 3), and some participants provided no or very few responses in one or more confidence categories across runs. This resulted in insufficient trial numbers for reliable condition-wise analyses. Therefore, we decided not to include the confidence ratings in the analysis.

Following fMRI acquisition, participants reported the extent to which they had used visual, spatial, tactile, and verbal learning strategies on a 5-point scale, with higher ratings indicating greater use of the respective strategy. Visual strategies were reported most strongly (mean = 3.0, SD = 1.0), followed by tactile (mean = 2.5, SD = 1.2), spatial (mean = 2.4, SD = 1.1), and verbal strategies (mean = 2.2, SD = 1.6). Open-ended responses indicated that participants frequently visualized the stimulus shapes (29.5%), assigned verbal labels to patterns (20.5%), focused on distinctive stimulus features such as movement direction (20.5%), associated stimuli with letters or numbers (18.2%), or concentrated on the tactile sensation itself (11.4%).

**TABLE 1.**
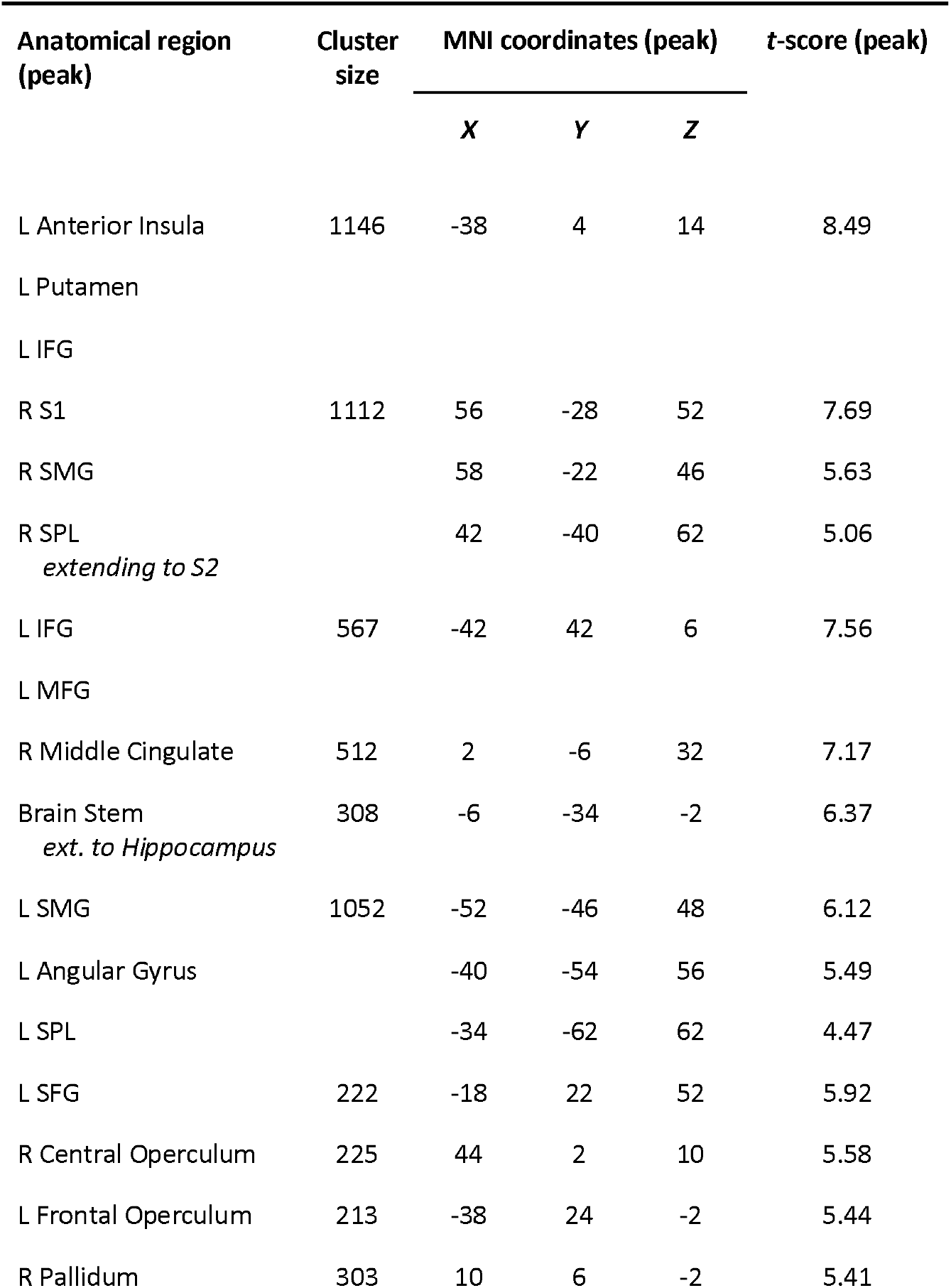
Brain regions showing greater activation during successful tactile memory retrieval (Hit > CR). Reported are the anatomical location of the peak voxel, cluster size (voxels), MNI coordinates of the peak voxel, and corresponding peak *t*-value. Results are shown at a voxel-wise threshold of *p* < 0.001, with cluster-level FWE correction at *p* < 0.05.

| Anatomical region<br>(peak) | Cluster<br>size | MNI coordinates (peak) |  |  | t-score (peak) |
| --- | --- | --- | --- | --- | --- |
|  |  | X | Y | Z |  |
| L Anterior Insula | 1146 | -38 | 4 | 14 | 8.49 |
| L Putamen |  |  |  |  |  |
| L IFG |  |  |  |  |  |
| R S1 | 1112 | 56 | -28 | 52 | 7.69 |
| R SMG |  | 58 | -22 | 46 | 5.63 |
| R SPL<br>*extending to S2* |  | 42 | -40 | 62 | 5.06 |
| L IFG | 567 | -42 | 42 | 6 | 7.56 |
| L MFG |  |  |  |  |  |
| R Middle Cingulate | 512 | 2 | -6 | 32 | 7.17 |
| Brain Stem<br>*ext. to Hippocampus* | 308 | -6 | -34 | -2 | 6.37 |
| L SMG | 1052 | -52 | -46 | 48 | 6.12 |
| L Angular Gyrus |  | -40 | -54 | 56 | 5.49 |
| L SPL |  | -34 | -62 | 62 | 4.47 |
| L SFG | 222 | -18 | 22 | 52 | 5.92 |
| R Central Operculum | 225 | 44 | 2 | 10 | 5.58 |
| L Frontal Operculum | 213 | -38 | 24 | -2 | 5.44 |
| R Pallidum | 303 | 10 | 6 | -2 | 5.41 |

### 3.2 Somatosensory cortex recruitment during tactile long-term memory recognition

To identify brain regions involved in successful tactile memory recognition, we contrasted trials in which learned stimuli were correctly recognized (Hits) with trials in which novel stimuli were correctly rejected (CRs). Importantly, the first-level model accounted for trial-by-trial differences in RT as well as activity associated with each of the four physical response buttons. The Hit > CR contrast revealed significantly greater activation in right S1, contralateral to the stimulated hand, extending into the supramarginal gyrus (SMG) and superior parietal lobule (SPL) (Figure 3; Table 1). Activation in right S2 was also evident within this cluster. Additional significant clusters included the left anterior insula extending into the pallidum and inferior frontal gyrus (IFG), left IFG extending into the middle frontal gyrus (MFG), left SMG extending into the angular gyrus and SPL, left superior frontal gyrus (SFG), bilateral frontal/central operculum, right pallidum, middle cingulate cortex (MCC), and the left hippocampus.

Hippocampal activation was predominantly observed in the left hemisphere. Inspection of individual functional data revealed susceptibility-related signal loss affecting parts of the right hippocampus in a subset of participants (Supplementary Figure 4), which may have reduced sensitivity to hippocampal activation in the right hemisphere.

**FIGURE 3.**
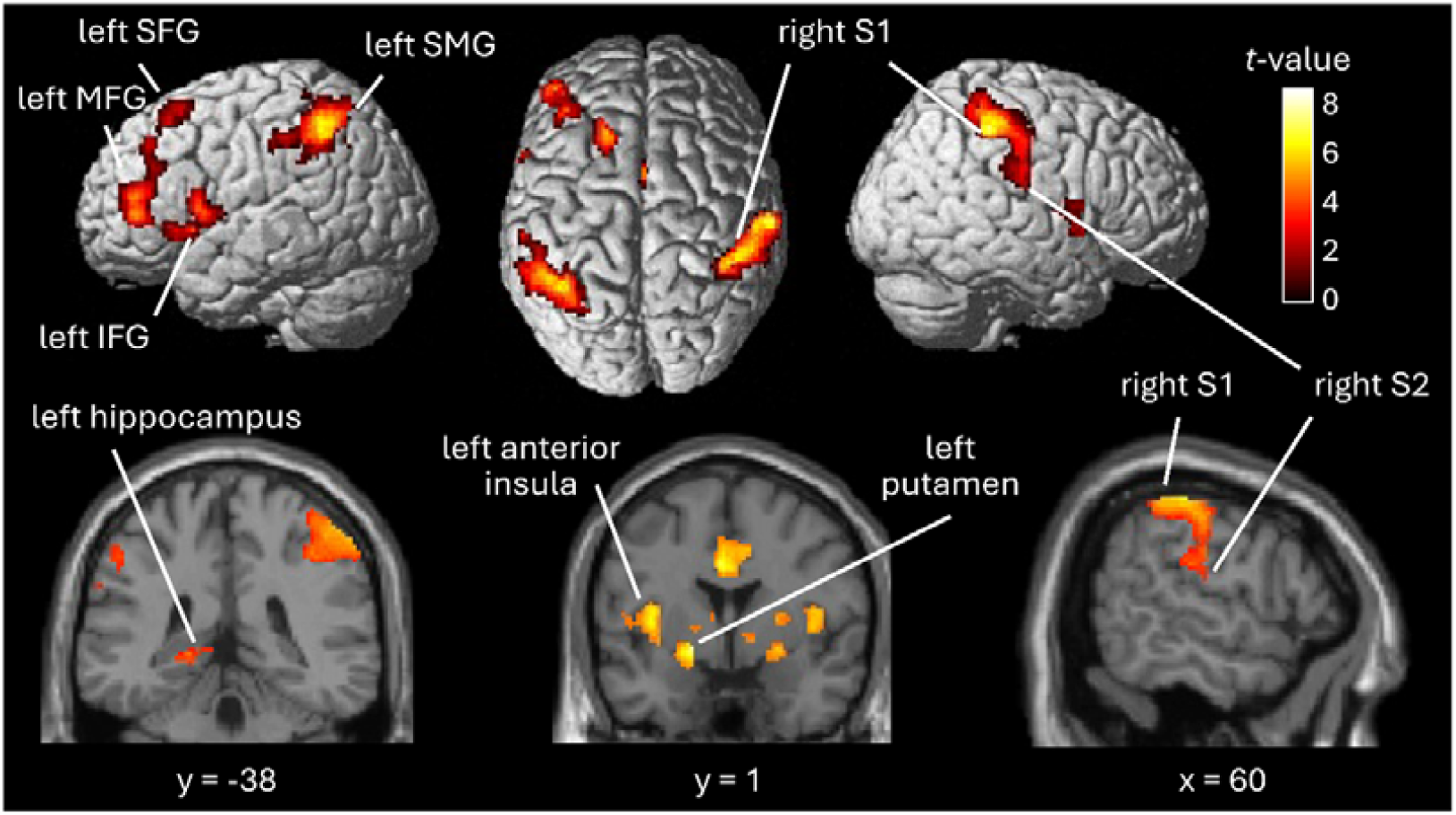
BOLD activation associated with successful tactile long-term memory recognition. Group-level activation for Hits > CRs, comparing successful recognition of previously learned tactile stimuli with correct rejection of novel stimuli are displayed. Surface renderings show the distributed cortical activation pattern, including contralateral right S1 and S2 and left-lateralized frontal and parietal regions. Coronal and sagittal sections highlight activation in the left hippocampus, left anterior insula, left putamen and right somatosensory cortex, respectively. Statistical maps are shown using a voxel-wise cluster-defining threshold of *p* < 0.001 (uncorrected), with significant clusters defined at *p* < 0.05, FWE-corrected at the cluster level. Peak coordinates, cluster sizes, and *t*-values are reported in Table 1.

To assess whether the observed activation pattern was influenced by individual differences in learning strategies, we repeated the second-level analysis including visual, spatial, tactile, and verbal strategy ratings as covariates. This analysis revealed a highly similar activation pattern, indicating that the principal effects were robust to individual differences in reported learning strategies (Supplementary Figure 5).

To further characterize the effects underlying the primary Hits > CRs comparison, we examined several supplementary contrasts. CRs > Hits, Hits > FAs, and Misses > CRs did not reveal any clusters meeting the predefined cluster-level FWE-corrected significance threshold. In contrast, Hits > Misses, directly comparing successful and unsuccessful recognition of previously learned stimuli, revealed greater activity in contralateral right S1 and left-lateralized frontoparietal regions, including the SMG, SFG, and MFG (Supplementary Figure 6). The S1 effect indicates that its recruitment was associated with successful recognition rather than prior stimulus exposure alone.

Together, these findings indicate that tactile memory retrieval engages both modality-specific somatosensory regions and distributed memory- and control-related networks, with supplementary analyses further linking contralateral S1 recruitment to successful recognition rather than prior stimulus exposure alone.

### 3.3 Hippocampal–somatosensory coupling during successful tactile long-term memory recognition

To characterize interactions between hippocampal memory systems and somatosensory cortex during tactile memory retrieval, we performed a PPI analysis using participant-specific seeds centered on local maxima within the left hippocampal activation identified for Hits > CRs. The PPI model accounted for trial-by-trial RT, activity associated with each of the four physical response buttons, and FA and Miss trials. Within an anatomical right S1/S2 mask derived from the JuBrain Atlas, Hits > CRs were associated with stronger functional coupling between the left hippocampus and the presumed hand area of contralateral right S1 and S2 (76 voxels; MNI: x = 58, y = −20, z = 52; *t* = 5.79) and S2 (8 voxels; MNI: x = 48, y = −32, z = 26; *t* = 3.97) at *p* < 0.001, uncorrected (Figure 4). Thus, successful recognition of learned tactile stimuli was associated with increased hippocampal coupling with somatosensory regions representing tactile information.

**FIGURE 4.**
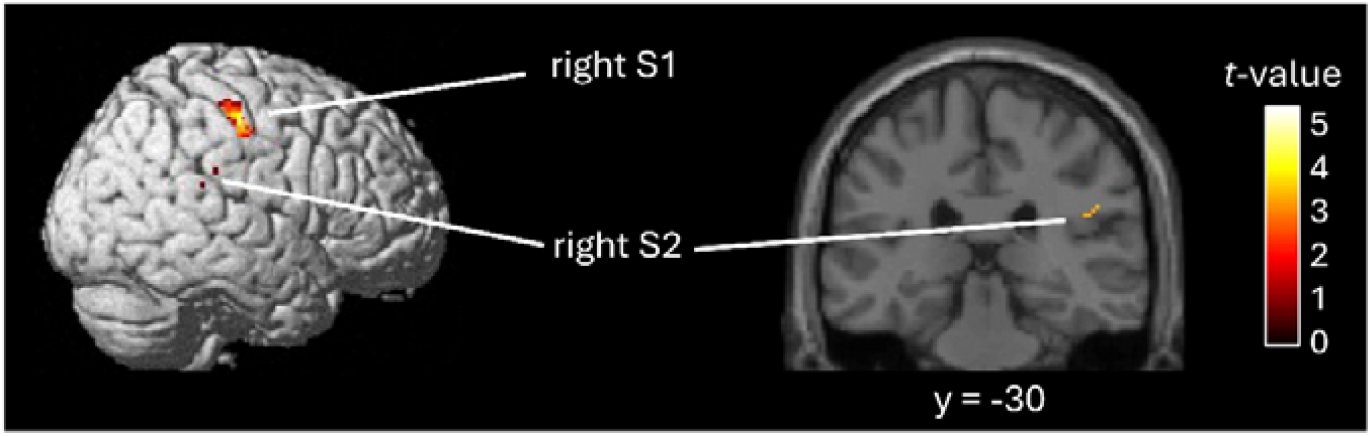
Hippocampal–somatosensory coupling during successful tactile long-term memory recognition. Group-level PPI analysis using the left hippocampus as the seed region identified regions showing stronger task-dependent coupling during Hits > CRs. The analysis was restricted to an anatomically defined right S1/S2 mask derived from the JuBrain Atlas and revealed increased hippocampal coupling with the presumed hand area of contralateral right S1 (76 voxels; MNI: x = 58, y = −20, z = 52; *t* = 5.79) and S2 (8 voxels; MNI: x = 48, y = −32, z = 26; *t* = 3.97). The coronal section highlights the S2 effect at y = −30. Results are shown at *p* < 0.001, uncorrected. The corresponding unmasked whole-brain PPI analysis is shown in Supplementary Figure 6.

An exploratory whole-brain analysis without the S1/S2 mask revealed a broader pattern of task-dependent hippocampal connectivity. Clusters in the left MFG and primary motor cortex (M1) reached whole-brain significance at *p* < 0.05, FEW-corrected at the cluster level, whereas other effects were observed only at the uncorrected threshold of *p* < 0.001 (Supplementary Figure 7).

## 4 Discussion

In this study, we investigated the neural correlates of tactile LTM using fMRI during recognition of learned dynamic spatiotactile stimulation events. In line with our hypotheses, successful recognition of learned stimuli, relative to correct rejection of novel stimuli, elicited increased activation in the contralateral hand representation of S1 as well as in contralateral S2, supporting a role of somatosensory cortices in tactile LTM representations. Successful recognition was additionally associated with activation in the left hippocampus. A PPI analysis seeded in this brain region further revealed increased retrieval-related functional coupling with both S1 and S2. Together, these findings suggest that tactile LTM retrieval involves coordinated interactions between hippocampal memory systems and modality-specific somatosensory cortices, consistent with the notion that S1 and S2 may contribute to distributed cortical memory engrams for tactile information.

A central unresolved question in tactile LTM is whether memory-related sensory representations are reinstated in S1, S2, or both. The involvement of somatosensory cortices during retrieval would be consistent with the concept of a distributed memory engram, according to which sensory cortices contribute modality-specific components of LTM representations that can be reactivated during retrieval (Josselyn & Tonegawa, 2020; Kitamura et al., 2017). Previous studies have provided partly divergent evidence: Bonda et al., (1996) implicated a ventral parieto-insular network including S2 during retrieval of haptically encoded information, whereas Stock et al., (2009) reported recruitment of S1 during haptic memory retrieval. Our findings reconcile these observations by showing that successful recognition of learned dynamic vibrotactile stimuli engages both contralateral S1 and S2. Activation in right S1 may reflect the reinstatement of relatively fine-grained, modality-specific sensory features of the learned stimulus representation. S1 is somatotopically organized (Sakai et al., 1995; Sanchez-Panchuelo et al., 2010) and retains detailed information about tactile input during WM (Schmidt et al., 2017), making it a plausible substrate for reactivation of sensory features established during learning. Consistent with this interpretation, the Hits > Misses contrast also revealed increased contralateral S1 activity when comparing successful and unsuccessful recognition of previously learned stimuli. Because both conditions comprised previously learned stimuli, this contrast controls for prior stimulus exposure and links greater S1 recruitment more specifically to successful retrieval. Moreover, the use of participant-specific stimulus sets minimized the possibility that this group-level effect was systematically driven by particular stimulus trajectories being both more memorable and elicited stronger S1 responses. This does not exclude sensory reinstatement as a mechanism; rather, it suggests that such reinstatement may be stronger or more effective when a tactile memory is successfully retrieved. Notably, successful recognition (Hits > CRs) also shows activation in the right SMG and SPL, suggesting recruitment of a broader somatosensory–parietal network during successful recognition. The SMG has been shown to contain stimulus-selective information during vibrotactile categorization (Malone et al., 2019), whereas the SPL has been implicated in tactile discrimination and the maintenance of tactile information (Kaas et al., 2013; Stoeckel et al., 2003). Together, this spatial pattern may reflect the recruitment of fine-grained sensory representations in S1 alongside higher-order parietal processing of tactile information during successful memory retrieval.

S2 may contribute at a complementary level of the tactile processing hierarchy. Compared with S1, S2 has been associated with bilateral integration, higher-order tactile processing, and more integrated somatosensory representations, properties that may support the integration of sensory information into mnemonic representations (Burton & Sinclair, 2000; Romo et al., 2012). This may be particularly relevant for the spatially dynamic nature of the stimuli used in the present study. Rather than presenting isolated, static tactile inputs, we deliberately designed each stimulus as a continuous tactile event with a defined beginning and end, thereby capturing an important feature of natural sensory experience while retaining precise experimental control. Recruitment of S2 during Hits > CRs is consistent with the ventral parieto-insular activation reported during haptic memory retrieval by Bonda et al., (1996) and suggests that successful tactile recognition engages multiple levels of the somatosensory hierarchy, with S1 contributing more fine-grained sensory information and S2 supporting more integrated spatiotemporal representations.

The anterior insula activation observed in our study further parallels the ventral parieto-insular involvement reported by Bonda et al., (1996). Notably, this cluster extended into the putamen, suggesting that additional components of this network may contribute to processing of dynamic tactile information during memory retrieval. The putamen has been implicated in temporal processing and motor timing (Merchant et al., 2015), as well as in the discrimination of moving tactile stimuli in non-human primates (Merchant et al., 1997) and humans (Numminen et al., 2004). Activity of the putamen has also been linked to the representation and retrieval of temporal stimulus information (Coull et al., 2008). Given the sequential nature of our tactile stimuli, its recruitment may therefore relate to the temporal organization of the learned tactile patterns.

Beyond these modality-specific effects, successful recognition recruited regions associated with broader episodic retrieval and cognitive control processes. Hippocampal recruitment is consistent with its well-established role in episodic retrieval and pattern completion, through which partial sensory input can trigger the reactivation of distributed cortical representations associated with a previous experience (Norman & O’Reilly, 2003; Rugg & Vilberg, 2013). The spatiotemporally extended nature of the stimuli is likely to have contributed to hippocampal involvement. As discussed by Dudai et al., (2015), temporally bounded experiences can be treated as mnemonic events, for which the hippocampal system binds information unfolding over time into a coherent episodic representation. Our dynamic tactile stimuli may therefore engage memory processes that extend beyond declarative memory for individual stimulus features, requiring the integration of successive sensory information into an event-like representation. In this sense, the paradigm provides a controlled approximation of the temporally extended experiences that episodic memory systems evolved to encode. Hippocampal activation was predominantly observed in the left hemisphere. However, susceptibility-related signal loss affected parts of the right hippocampus in a subset of participants (Supplementary Figure 4), potentially reducing sensitivity and limiting conclusions about the hemispheric distribution of hippocampal involvement.

Successful recognition (Hits > CRs) additionally recruited an extensive parietal network, including bilateral SMG and left angular gyrus and SPL. The angular gyrus is consistently implicated in successful episodic retrieval and has been proposed to integrate retrieved features into a coherent mnemonic representation and contribute to the subjective experience of recollection (Cabeza et al., 2008; Ramanan et al., 2018). More broadly, lateral parietal cortex has been proposed to mediate attentional interactions with retrieved memory contents, with ventral parietal regions particularly responsive when mnemonic information is successfully recovered (Cabeza et al., 2008; Ciaramelli et al., 2010). The bilateral parietal recruitment observed here may therefore reflect higher-order integration of retrieved tactile information with mnemonic evidence during successful recognition.

Frontal recruitment encompassed left IFG and MFG, SFG, as well as bilateral frontal/central opercular regions. Prefrontal regions are consistently recruited during recognition memory and have been associated with controlled retrieval, selection among competing representations, and post-retrieval monitoring and evaluation (Henson et al., 1999; Rugg & Wilding, 2000). Such processes are particularly relevant to the present task, in which participants had to evaluate an incoming tactile sequence against previously learned representations and classify it as learned or novel. The accompanying middle cingulate activation may further reflect cognitive and response-related control processes engaged during this decision (Vogt, 2016). Thus, rather than representing tactile memory content itself, frontal and cingulate recruitment may reflect domain-general control processes supporting successful recognition. Together, these findings suggest that tactile long-term memory retrieval recruits the hippocampus alongside distributed parietal and frontal systems, complementing the modality-specific representations observed in somatosensory cortex.

The regional activation findings were complemented by evidence for retrieval-dependent interactions between hippocampal and somatosensory systems. PPI analyses demonstrated that successful retrieval was accompanied by increased task-dependent coupling between the left hippocampus and both contralateral S1 and S2 compared with correct rejection of novel stimuli. This finding complements the regional activation effects by showing that successful recognition was associated not only with recruitment of hippocampal and somatosensory regions, but also with stronger functional interactions between them. Hippocampal–neocortical interactions are central to LTM formation and retrieval: the hippocampus rapidly binds distributed aspects of an experience, while their repeated coordinated reactivation contributes to the stabilization of distributed cortical memory representations (Takehara-Nishiuchi, 2021; Wang & Morris, 2010). During subsequent retrieval, partial sensory cues can engage hippocampal pattern-completion mechanisms that facilitate reinstatement of associated cortical representations (Norman & O’Reilly, 2003; Rugg & Vilberg, 2013; Staresina & Wimber, 2019). The increased hippocampal coupling with S1 and S2 during successful tactile recognition is consistent with such retrieval-dependent coordination between hippocampal memory processes and modality-specific cortical representations. Although PPI does not establish the direction of information flow or demonstrate reinstatement itself, the convergence of regional activation and coupling effects supports the involvement of somatosensory cortex within a distributed hippocampal-cortical retrieval network.

Several behavioral and control analyses further inform the interpretation of these neural effects. Recognition performance increased significantly across the three learning sessions, demonstrating successful acquisition of the tactile stimuli, and remained stable across the six fMRI runs, with an overall recognition accuracy of 62.2%. Although performance was above chance, the moderate accuracy underscores the difficulty of learning and discriminating the 16 complex, tactile stimuli. Importantly, RTs differed significantly across recognition outcomes, with responses fastest for Hits and slowest for Misses. Because differences in response latency could otherwise contribute to the neural differences between recognition conditions, trial-by-trial RT was explicitly modeled as a common parametric nuisance regressor in the first-level GLM. Moreover, the four physical response buttons were modeled separately at the time of the button press. Thus, the principal Hits > CRs effects were estimated after accounting for variance associated with both RT and the specific motor response. The comparable distribution of physical button presses across recognition outcomes further argues against systematic differences in button use as an explanation for the observed effects.

Participants nevertheless differed considerably in how they approached learning the tactile stimuli. Even though they were instructed to focus on the tactile properties of the stimuli for memorization, they indicated in the debriefing questionnaire handed out after scanning that they mostly used a visual strategy to memorize the patterns, followed by spatial, tactile, and verbalized strategies. The influences of visual imagery processes in nonvisual recognition paradigms are often reported (Sathian, 2016) and thus, it is not surprising that these processes were probably involved in the current study. However, including participants’ reported visual, spatial, tactile, and verbal learning strategy ratings as covariates in the second-level GLM yielded a highly similar activation pattern, indicating that the observed retrieval network was robust to individual differences in reported learning strategies. Although visual imagery may have contributed to task performance, it is therefore unlikely to account for the principal activation and connectivity patterns reported here.

Several limitations define directions for future work. Confidence responses were unevenly distributed across participants, precluding reliable separation of recognition success from subjective confidence, which is known to modulate activity during episodic recognition (Henson et al., 2000). The spatial resolution of the present data also precluded characterization of memory representations at finer anatomical scales. High-field submillimeter fMRI could test whether retrieval reinstates digit-specific representations within S1 and whether S1 and S2 encode complementary features of remembered tactile stimuli (Sanchez-Panchuelo et al., 2010). Depth-resolved measurements could further examine whether retrieval-related activity shows laminar profiles consistent with top-down feedback, exploiting the differential laminar organization of feedforward and feedback signals (Lawrence et al., 2019; Self et al., 2019). In parallel, high-resolution imaging of hippocampal subfields could test whether DG/CA3 and CA1 make dissociable contributions to tactile retrieval (Lazarova et al., 2026). Combining these approaches could reveal how hippocampal computations interact with fine-scale somatosensory representations during tactile memory retrieval.

Finally, tactile memory may have translational relevance. Intrusive memories in PTSD are often dominated by vivid sensory-perceptual components, and altered sensory processing is increasingly recognized as part of PTSD pathophysiology (Ehlers et al., 2004; Fleming et al., 2024) (Ehlers et al., 2004; Cushing et al., 2023). Extending tactile memory paradigms to clinical populations could therefore test whether intrusive memories involve altered reinstatement of somatosensory representations or altered hippocampal–somatosensory interactions.

In conclusion, our findings suggest that tactile LTM retrieval engages multiple levels of the somatosensory hierarchy: S1 may reinstate fine-grained, somatotopically organized sensory information, whereas S2 may integrate higher-order spatiotemporal stimulus features. Increased coupling of both regions with the hippocampus further links these modality-specific representations to hippocampal retrieval mechanisms, embedded within a broader frontoparietal network supporting retrieval and recognition decisions. Together, these findings extend distributed models of long-term memory to the tactile domain and provide a framework for investigating how sensory memory representations are altered in clinical conditions affecting memory and somatosensory processing.

## 5 Acknowledgements

We thank Esther Kuehn for her helpful discussions and Christian Kainz for his support with data acquisition and COVID-19 safety measures during the study.

## Notes

**Conflict of Interest Disclosure** The authors declare no conflicts of interest.

### Competing Interest Statement

The authors have declared no competing interest.

