## Supplementary figures for "Remembering touch: Somatosensory cortex supports long-term memory for dynamic vibrotactile patterns"

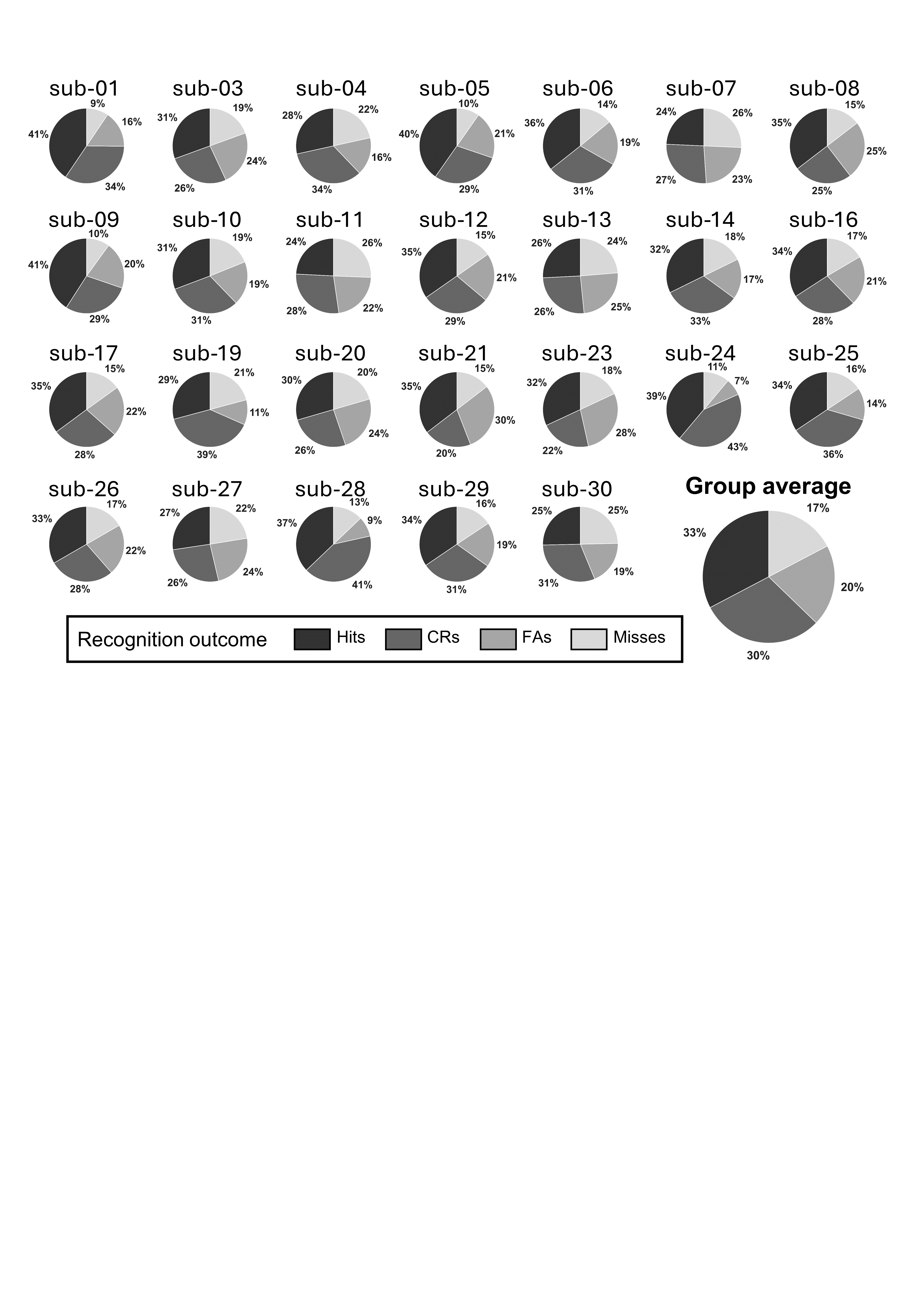

**SUPPLEMENTARY FIGURE 1**. Distribution of recognition outcomes during the fMRI recognition task.

Pie charts show the proportion of trials classified as Hits, Correct Rejections (CRs), False Alarms (FAs), and Misses for each of the 26 participants included in the fMRI analyses. The larger pie chart shows the mean proportion of each response category across participants.

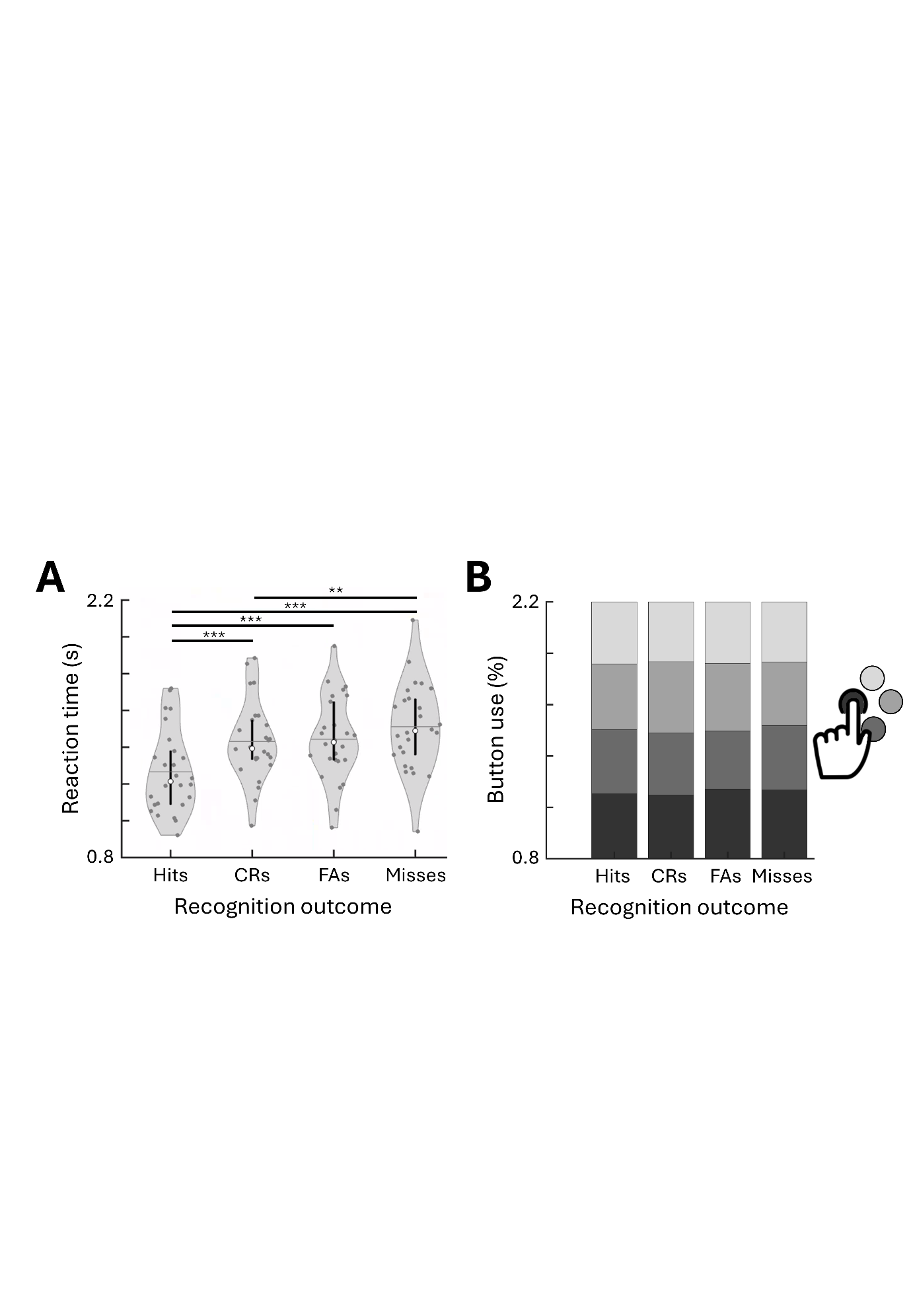

**SUPPLEMENTARY FIGURE 2**. Response times and response-button use across recognition outcomes.

(A) Distribution of reaction times for Hits, Correct Rejections (CRs), False Alarms (FAs), and Misses across participants. Gray dots indicate individual participants mean response times, and violin widths represent the kernel-density estimate. White circles indicate the median, vertical black lines span the interquartile range (25th–75th percentiles), and horizontal lines indicate the mean. Asterisks indicate significant Bonferroni-corrected pairwise comparisons following the repeated-measures ANOVA (**: *p* < 0.01, ***: *p* < 0.001). (B) Relative use of the four response buttons for each recognition outcome. Gray shades correspond to the four physical button positions indicated by the button box schematic. Response-button mapping was randomized across trials; no coincidental bias is indicated.

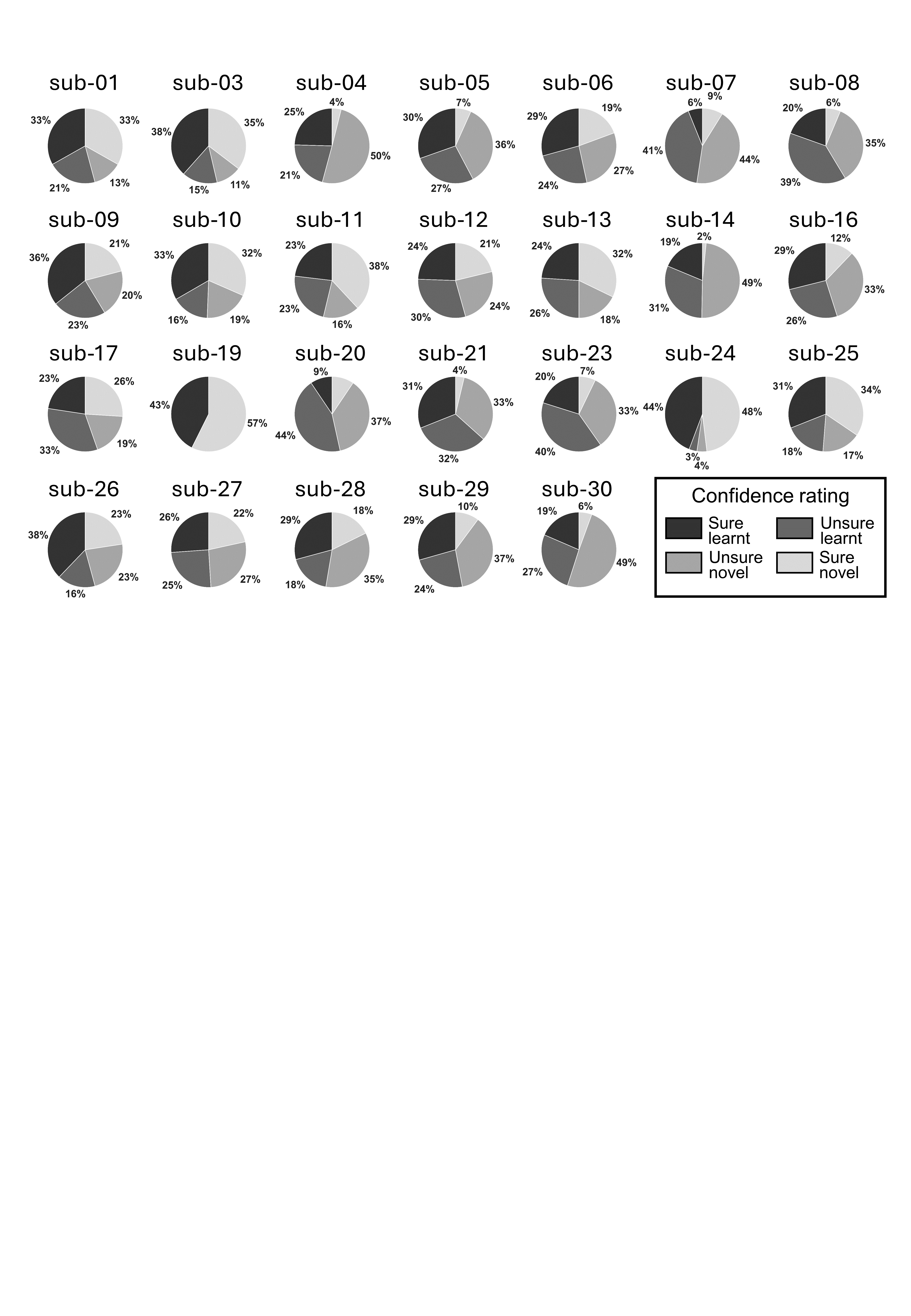

**SUPPLEMENTARY FIGURE 3**. Distribution of confidence ratings across participants.

For each participant, the relative proportion of responses categorized as ‘sure learnt’, ‘unsure learnt’, ‘sure novel’, and ‘unsure novel’ across all trials during the fMRI session is displayed. Whereas a small number of participants showed relatively balanced response distributions (e.g., sub-12, sub-17, sub-27, sub-28), most participants exhibited strongly unbalanced response patterns with underrepresented or absent categories (e.g., sub-04, sub-07, sub-08, sub-14, sub-19, sub-20, sub-21, sub-23, sub-24, sub-30). Due to this uneven distribution of confidence ratings, no confidence-based analyses were performed for the fMRI data.

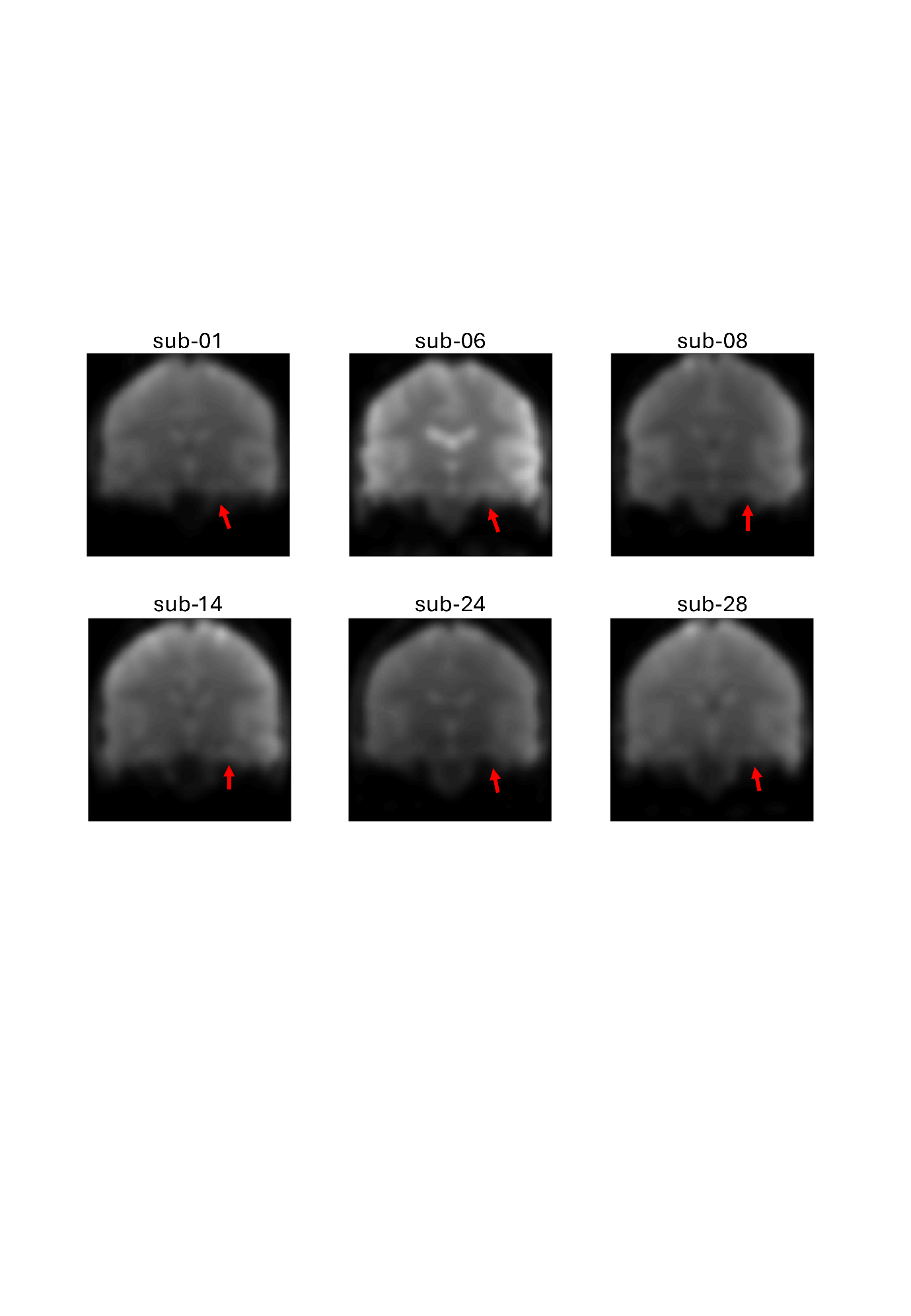

**SUPPLEMENTARY FIGURE 4**. Susceptibility-related signal loss in the right hippocampus.

Representative coronal functional images from the first run of six participants illustrating susceptibility-related signal loss affecting portions of the right hippocampus. Red arrows indicate regions of reduced signal in the medial temporal lobe encompassing parts of the right hippocampus. Reduced signal in this region limited sensitivity to right hippocampal activation and therefore precluded conclusions about the hemispheric distribution of hippocampal involvement.

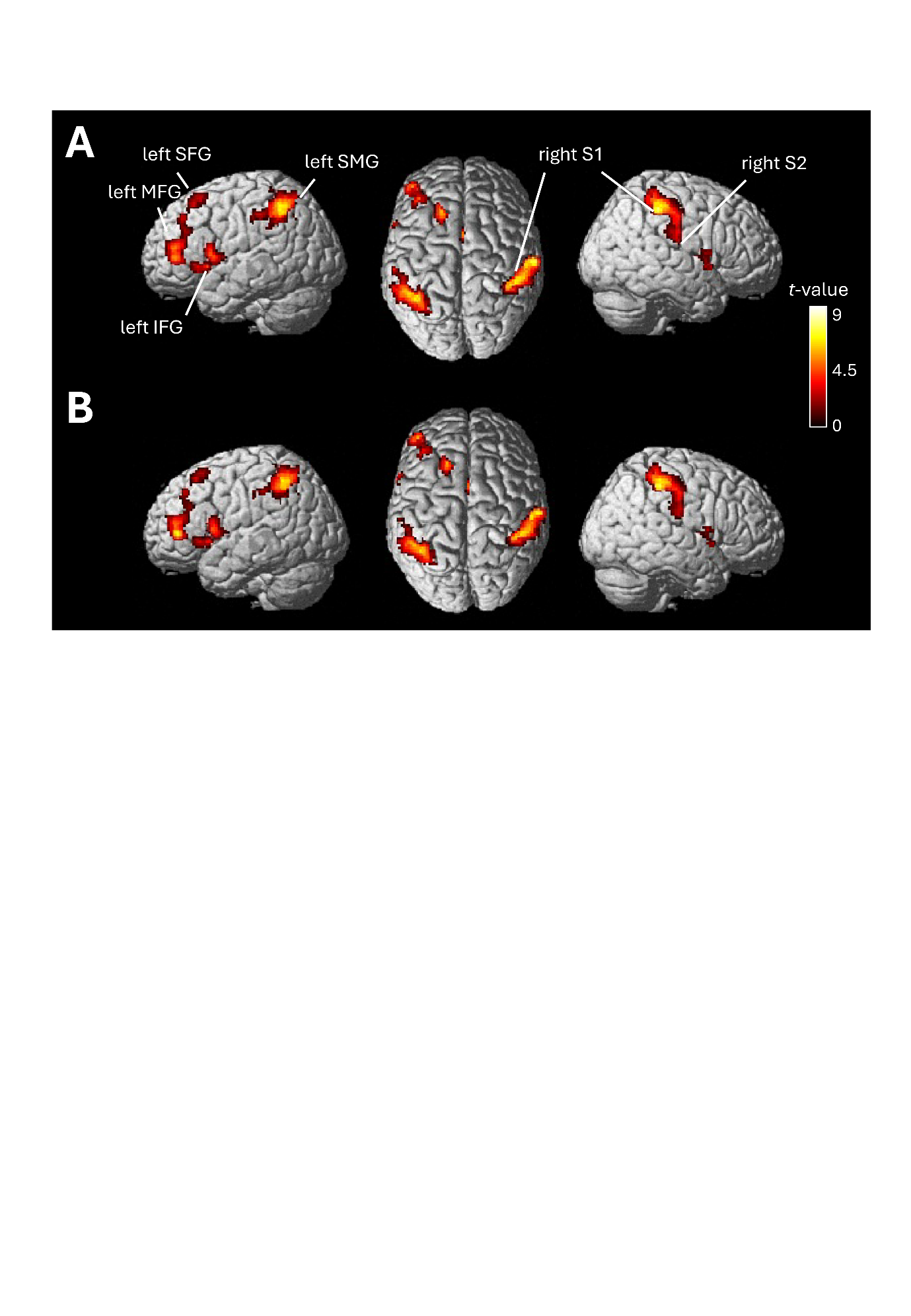

**SUPPLEMENTARY FIGURE 5.** Robustness of tactile memory-related activation to individual learning strategies.

(A) Group-level activation for the Hits > CRs contrast from the primary analysis, also presented in Figure 3. (B) The Hits > CRs analysis was repeated with participants’ self-reported visual, spatial, tactile, and verbal learning-strategy ratings from the post-scan debriefing questionnaire entered as mean-centered covariates of no interest in the second-level model. The highly similar activation patterns indicate that the principal effects associated with successful tactile recognition were robust to individual differences in reported learning strategies. Statistical maps are shown using a voxel-wise cluster-defining threshold of *p* < 0.001 (uncorrected), with significant clusters defined at *p* < 0.05, FWE-corrected at the cluster level. S1, primary somatosensory cortex; S2, secondary somatosensory cortex; IFG, inferior frontal gyrus; MFG, middle frontal gyrus; SFG, superior frontal gyrus; SMG, supramarginal gyrus.

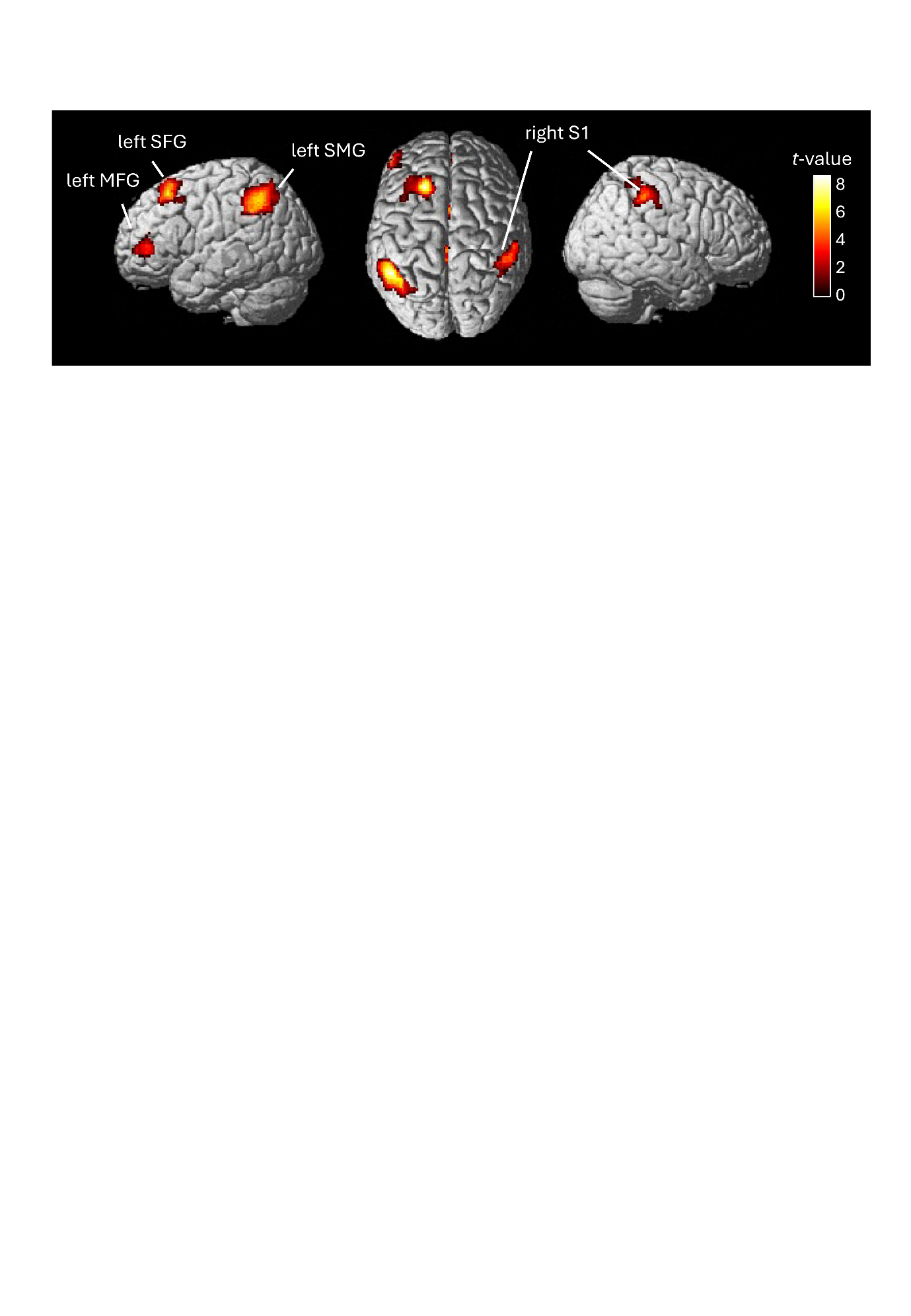

**SUPPLEMENTARY FIGURE 6**. Neural activity associated with successful recognition of learned tactile stimuli.

Group-level activation for the Hits > Misses contrast, comparing successfully and unsuccessfully recognized previously learned stimuli. As both conditions comprised previously learned stimuli, the contrast isolates differences associated with recognition success. Greater activation for Hits was observed in contralateral right primary somatosensory cortex (S1) and left-lateralized frontoparietal regions, including the supramarginal gyrus (SMG), superior frontal gyrus (SFG), and middle frontal gyrus (MFG). Statistical maps are shown using a voxel-wise cluster-defining threshold of *p* < 0.001 (uncorrected), with significant clusters defined at *p* < 0.05, FWE-corrected at the cluster level.

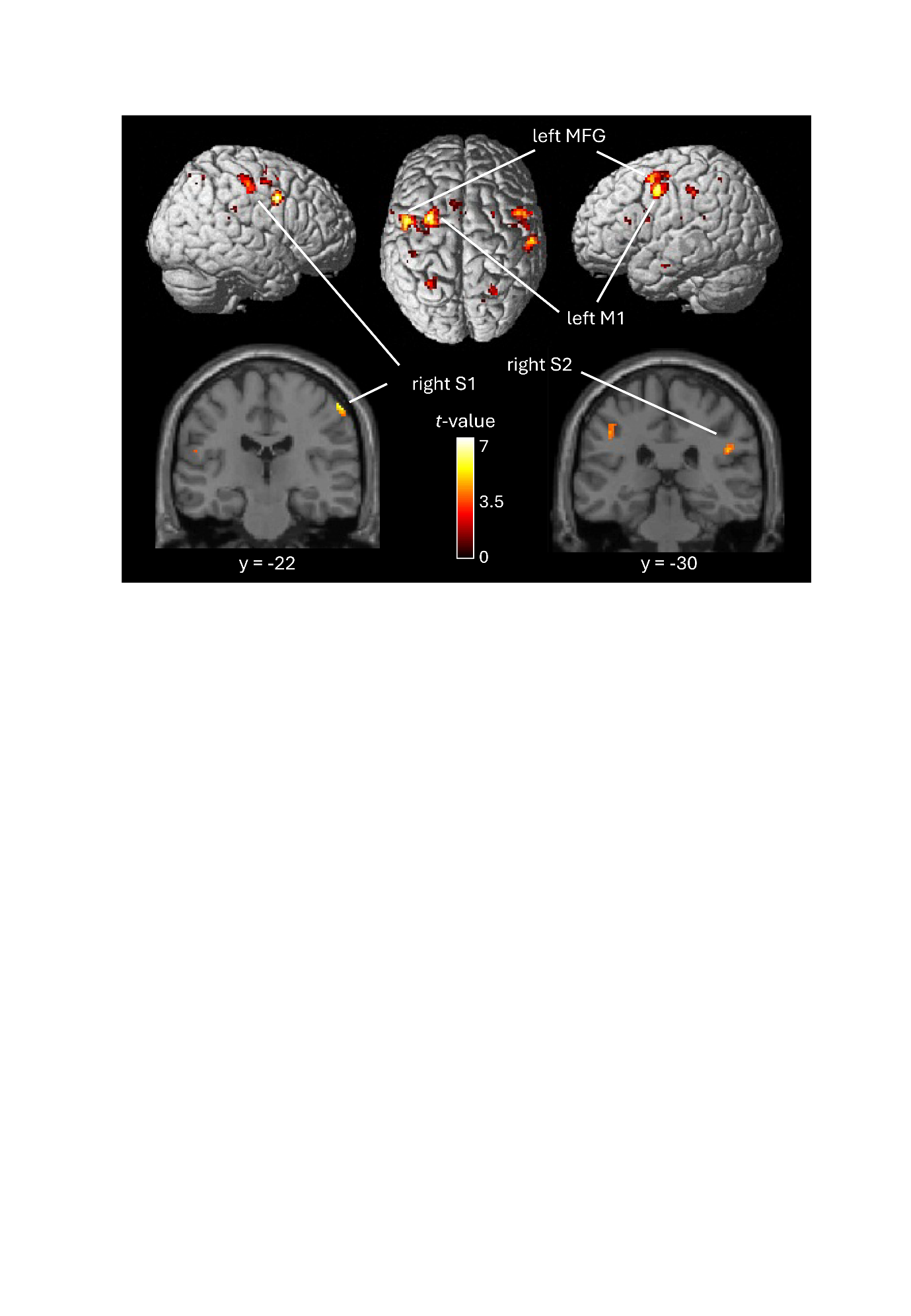

**SUPPLEMENTARY FIGURE 7**. Whole-brain hippocampal functional coupling during tactile long-term memory recognition.

Unmasked whole-brain PPI analysis showing regions with stronger task-dependent functional coupling with the left hippocampus during Hits relative to CRs. Clusters in the left middle frontal gyrus (MFG) and primary motor cortex (M1) reached significance at *p* < 0.05, FWE-corrected at the cluster level. Additional effects were observed at p < 0.001, uncorrected, including in contralateral right primary (S1) and secondary (S2) somatosensory cortex. Coronal sections illustrate the effects in right S1 (y = −22) and right S2 (y = −30). Results are displayed at *p* < 0.001, uncorrected.
